# A semi-automatic workflow to process camera trap images in R

**DOI:** 10.1101/2022.10.05.510927

**Authors:** Hanna Böhner, Eivind Flittie Kleiven, Rolf Anker Ims, Eeva M. Soininen

## Abstract

Camera traps have become popular for monitoring biodiversity and animal populations. Artificial intelligence is increasingly used to automatically classify large image data sets produced by camera traps and many tools that incorporate machine-learning models for automatic image classification have been developed over the last years. However, it is still challenging to combine tools for automatic classification with other tools for processing camera trap images and to adapt these tools to a specific study. Therefore, we propose a semi-automatic workflow for processing camera trap images in R. The workflow includes managing raw images, automatic image classification, a quality check of automatic image labels as well as the possibilities to retrain the model with new images and to manually review subsets of images to correct image labels. We illustrate the workflow with a case-study from the small mammal monitoring program of the Climate-ecological Observatory for Arctic Tundra. We first trained a classification model for small mammals and then transferred the model to new images, including images from newly established camera traps. We could show that retraining the original model with a small number of new images increased model performance and therefore highlight the importance of verifying automatic image labels when a model was transferred to new images. Furthermore, retraining the original model also decreased the time needed for manually reviewing images and correcting image labels substantially. Thus, the proposed workflow results in a data set with high accuracy and minimizes time needed for labeling images manually. This is especially useful for long-term monitoring where new images have to be processed continuously and methods have to be adapted over time. We provide all R scripts and the classification model for small mammals to make the workflow accessible to other ecologists.

## 1. Introduction

The increasing use of automated sensor networks such as wildlife camera traps, phenology cameras and acoustic sensors has advanced ecological research by providing high-resolution and large-scale data for better understanding of ecological processes and improving ecological forecasting (Farley et al., 2018). These sensor networks usually produce enormous amounts of data which need to be stored and processed, an often challenging task for ecologists. For example, camera traps which have become widely used tools for wildlife monitoring in the last decades (Burton et al., 2015; Steenweg et al., 2017) can easily accumulate thousands of images in a short time. Traditionally, the images are reviewed by humans who manually extracted data such as the presence of a species or the number of individuals on the image. This is a very time consuming process and has limited the use of camera trap data so far (Glover-Kapfer et al., 2019).

To overcome the challenge of manually processing huge amounts of data, artificial intelligence is increasingly used in ecology, for example to automate the classification of camera trap images (Christin et al., 2019). Deep neural networks are a type of machine learning models that have proven to be especially useful for image recognition (Krizhevsky et al., 2012). Several researchers developed neural networks for animal detection and classification on camera trap images with great success (e.g. Norouzzadeh et al., 2018; Tabak et al., 2019; Willi et al., 2019; Zualkernan et al., 2022). These models can for example separate empty images from images containing an animal (e.g. Beery et al., 2019), identify species, count individuals, and categorize animal behaviour with an accuracy that matches or even outperforms humans (Norouzzadeh et al., 2018). However, developing and training neural networks requires programming skills and have therefore often been tasks for computer scientists (Tabak et al., 2020). To facilitate the use of machine learning in ecology, several neural networks trained on large image data sets are now publicly available via simple user interfaces. For example, Tabak et al. (2020) developed the R-package ‘Machine Learning for Wildlife Image Classification (MLWIC2), which provides functions for classifying new camera trap images with the provided models and for training new classification model using own images. Besides tools for automatic image classification, a range of software tools for camera trap data management and image annotation have been developed since the use of camera traps became popular (see review by Young et al., 2018). More recently, machine learning has been implemented in annotation programs to allow for semi-automatic workflows which combine automatic classification and human labeling. AIDE (Annotation Interface for Data-driven Ecology) is a open-source web framework for image annotation that also provides tools for training machine learning models (Kellenberger et al., 2020). Timelapse (http://saul.cpsc.ucalgary.ca/timelapse/) is an open-source annotation software that incorporates the output from a machine learning model and can be used to verify and correct model labels. However, Timelapse works only with the output from a specific detection model, the MegaDetector (Beery et al., 2019), and therefore can not be used if another model type was used for automatic classification.

Despite these recent advances, complete and flexible workflows for processing camera trap images from long-term monitoring programs are to our knowledge still rare. Besides automatic image classification, such workflows should include a quality check where the model performance is evaluated, followed by a step where automatic image labels can be reviewed and corrected manually. Although many tools for camera trap image processing are available already, they are not easily streamlined into a complete workflow because different programs and platforms have been developed for different tasks. In addition, these tools are often not very flexible and it might be difficult to adjust them to the the different needs of different camera studies. Due to these shortcomings, researchers have often rather developed their own camera data management tool than using available ones. Most of the avialable tools focus on either automatic or manual image classification, although a combination might often be most appropriate. In many cases, neural networks will rather accelerate manual classification than completely replace it (Vélez et al., 2022; Greenberg) because the transferability of neural networks to new images is known to be problematic (Norouzzadeh et al., 2018) and classification models usually perform better for some species than for others. Thus, the importance of a verifying automatic image labels has been emphasized (Christin et al., 2021). However, it is difficult to find examples for how such a quality check and possible correction of automatic image labels are carried out and incorporated in a workflow. This is especially important if camera traps are used in long-term monitoring programs where a model is trained with images taken over certain time period and then applied to classify new images taken after the model was trained. In this case, it will be important to verify that a previously trained model performs well for images from the same sites over time and for new sites which might be added to the monitoring program over time. Even species composition can change over time and it may therefore be necessary to adapt the model when conditions are changing.

In this study, we suggest a semi-automatic workflow for processing and classifying images from camera traps. The workflow includes organising and processing raw images retrieved from the camera traps, training and applying an automatic image classification model, a quality check of the automatic image labels ant the correction of wrong labels. In addition, we incorporate training a classification model and retraining the model with new images in the workflow. We demonstrate the workflow using images from a long-term small mammal monitoring program. Most publicly available classification models are trained with image data sets dominated by larger species, often ungulates, whereas images from small mammal cameras traps are dominated by small species such as voles or mustelids. Therefore, existing models can not be applied to these images and we provide a classification model for small mammals in the Fennoscandian tundra.

The complete workflow is performed in the statistical software R (R Core Team, 2022), a programming language used by most ecologists (Lai et al., 2019). Instead of a tool with a graphical user interface or an R package, we provide R scripts or Shiny apps for all steps of the workflow, including training a image classification model. R scripts are very flexible and can be easily adapted to fit other studies by ecologists with an average knowledge of R.

## 2. Description of the workflow

The semi-automatic workflow includes a processing pipeline from images downloaded in the field to a quality checked data set ready to be used in ecological analyses. In addition, we give an example for training an image classification model in R and for adapting the model to new data (Figure 1).

**Figure 1:**
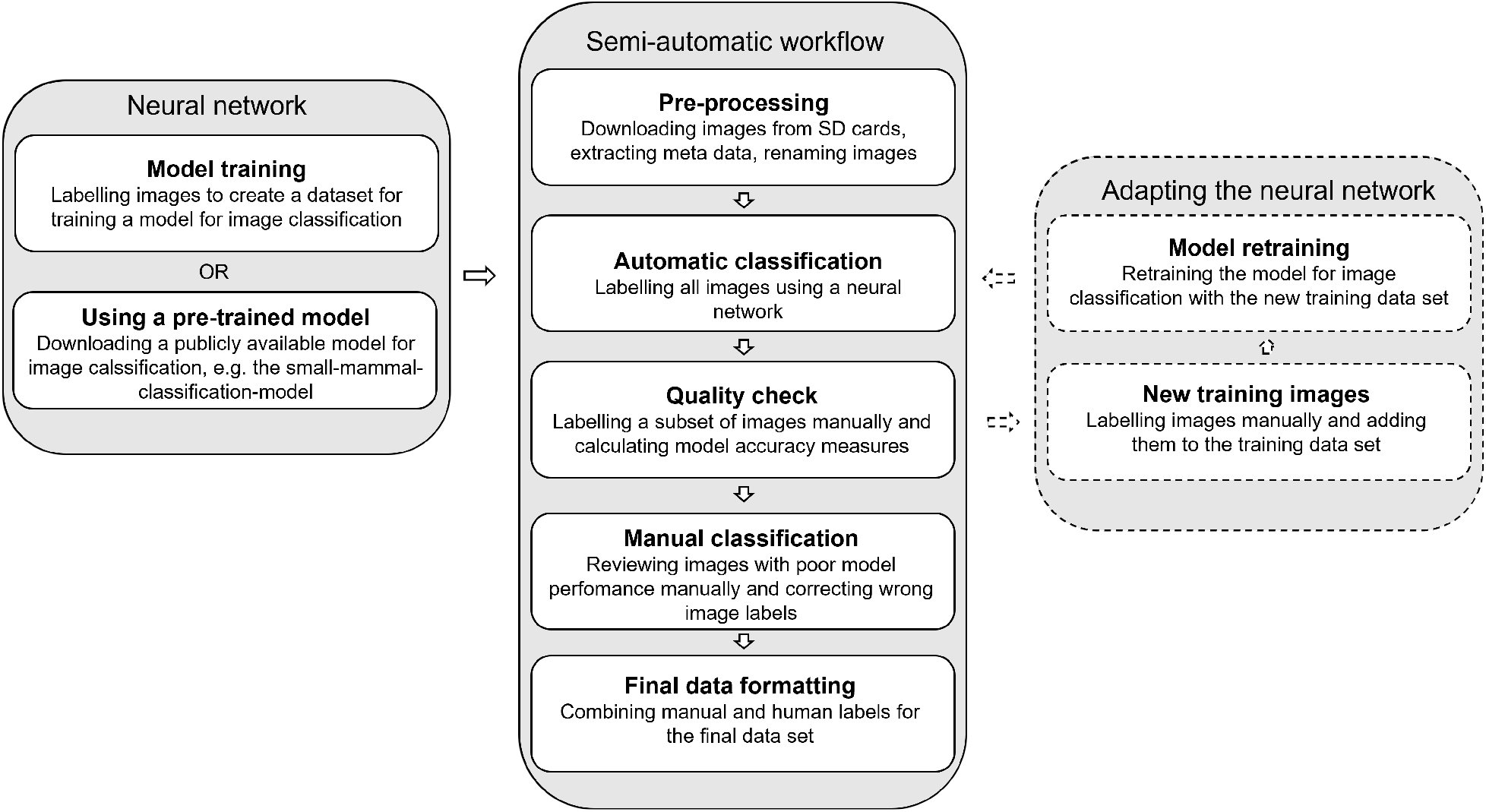
The steps of a semi-automatic workflow for classifying camera trap images.

### 2.1. Pre-processing

First, after retrieving the images from the cameras, we recommend renaming all images with a unique name. Camera trap images are usually automatically named with a generic name such as ‘IMG_0001.JPG’ by the camera and thus, images from different camera traps have the same names. This will cause confusion in large camera trap studies with many cameras running over a long time. Furthermore, image metadata saved with the image, such as date and time when the image was taken or temperature can be extracted and saved in a metadata-file within the same step.

### 2.2. Automatic classification

Then, all images are labeled automatically using an image classification model. The model can either be trained specifically for the data set or a pre-trained, publicly available model can be utilized. The output of classification models is a value between 0 and 1 per class, describing the confidence of the model that the image belongs to a certain class. The class with the highest confidence is usually extracted as the automatic image label.

### 2.3. Quality check

Image classification models often generalize poorly to new data, and thus, verification of the automatic image labels after classifying a new images is important. The quality check therefore includes labeling a subset of images manually and calculating accuracy measures such as prediction accuracy, precision, recall and F1 scores (Sokolova and Lapalme, 2009). We suggest a quality check in three steps that allows evaluation of (i) overall and (ii) per-class model performance as well as (iii) setting a confidence threshold. i) We recommend labeling a random subset of images manually for calculating overall model accuracy. ii) Since image data sets are often unbalanced, classes with few images might not be included in the random subset. In order to calculate per-class model accuracy, we recommend selecting random images of each class and labeling these manually in addition. iii) To determine a confidence threshold (confidence level above which model labels are deemed to be high accuracy and thus can be accepted), we recommend to select random images per confidence class for manual labeling, i.e. images classified with a confidence between 0.1 and 0.2, 0.2 and 0.3 and so on. Prediction accuracy can then be calculated for each confidence class.

### 2.4. Model retraining

Based on the quality check, the researcher can decide whether model performance is satisfying or if the model performance should be improved by selecting new training images and retraining the model. If images from new sites have been classified, model performance can be improved by including images from these sites in the training data set. If the model had problems with some species, it might help to include more images of these species in the training data set. When selecting new training images, the model output from the original model can be helpful to find images that meet a certain criteria, e.g. to find images of a certain species.

### 2.5. Manual classification

If the model performance is in general satisfying but poor for a specific type of images, for example for images with low confidence or for images of one species, the labels of these images should be reviewed manually and corrected if necessary.

### 2.6. Final data formatting

The automatic and manual image labels are combined in the final data set that then can be used in ecological analyses. The data set provides image metadata, animal observations (i.e. automatic and manual image labels) and image quality information.

## 3. Case study

In this example, we first trained a classification model using images that were collected over several years and labeled manually. Then, we used the model to automatically classify the images from a new year and from new camera trap types, performed a quality check, retrained the model with new images and corrected wrong automatic image labels. All R scripts needed for the workflow and detailed instructions for using the scripts and following the workflow are available on GitHub (https://github.com/hannaboe/camera_trap_workflow). We also provide the classification model for small mammals and our training data set which can be used to retrain the model together with additional images.

### 3.1. Image data set

We used images from long-term monitoring of small mammals from the the Climate-ecological Observatory for Arctic Tundra (Ims et al., 2013). The first camera trapping sites were established in 2014 on the Varanger peninsula in Northern Norway (Mölle et al., 2022). More traps were added in the following years, resulting in a total of 92 camera trap sites by 2020. The camera traps established during these years were located in two habitats (hummock tundra and snowbed habitat), and at two study areas (Komagdalen and Vestre Jakobselv) (Figure 2. The target species of the monitoring are Norwegian lemmings (*Lemmus lemmus*), voles (*Myodes rufocanus* and *Microtus oeconomus*) as well as their predators, stoats (*Mustela erminea*) and least weasels (*Mustela nivalis*). Furthermore, shrews (*Sorex ssp*.) and small birds are also frequently recorded.

**Figure 2:**
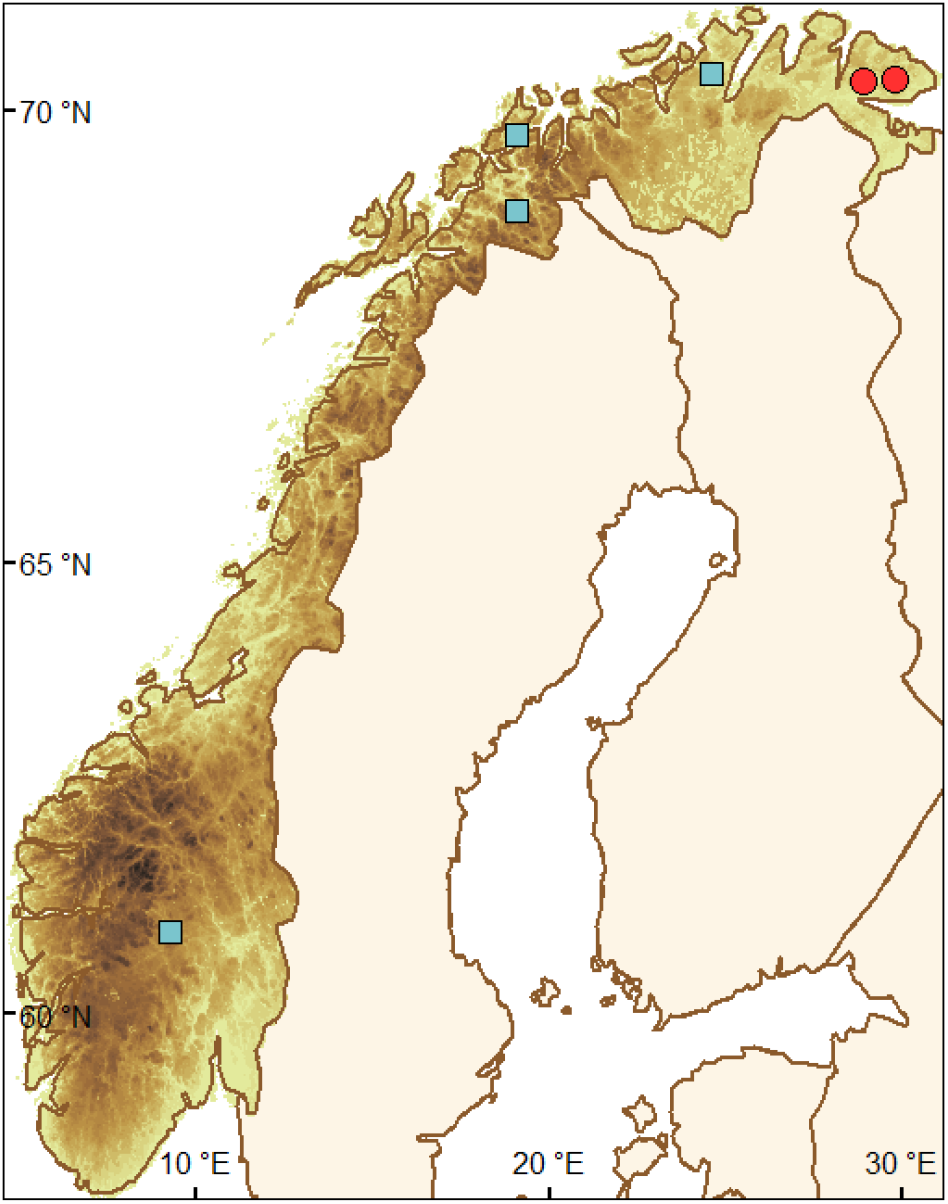
The location of the two camera trapping areas (Komagdalen and Vestre Jakobselv) of the small mammal long-term monitoring program on the Varanger peninsula are represented with red dots. The location of other small mammal camera trapping areas Valdres, Kirkesdalen, Håkøya, Porsanger are represented with blue squares. Images of these areas where used to extend the training data set.

The camera trap was developed by Soininen et al. (2015) and consisted of a Reconyx camera (Customized from Reconyx S750, Reconyx Inc., Holmen, WI, USA) mounted on the ceiling of a metal box that functions as a tunnel where small mammals can pass through (Figure 3). The cameras were programmed to take motion sensor images with two images per trigger and a quiet period of one minute between images. In addition, to monitor the functionality of the camera traps (battery life, intrusion with snow, ice, and water) two time-lapse images were taken per day. Most images are empty or contain a single animal on the floor of the box in a relatively fixed distance from the camera lens. However, animals have different positions and sometimes only parts of the animal is visible in the openings of the box. Snow, ice, water and vegetation can accumulate in the boxes and cause variable background (Figure 4).

**Figure 3:**
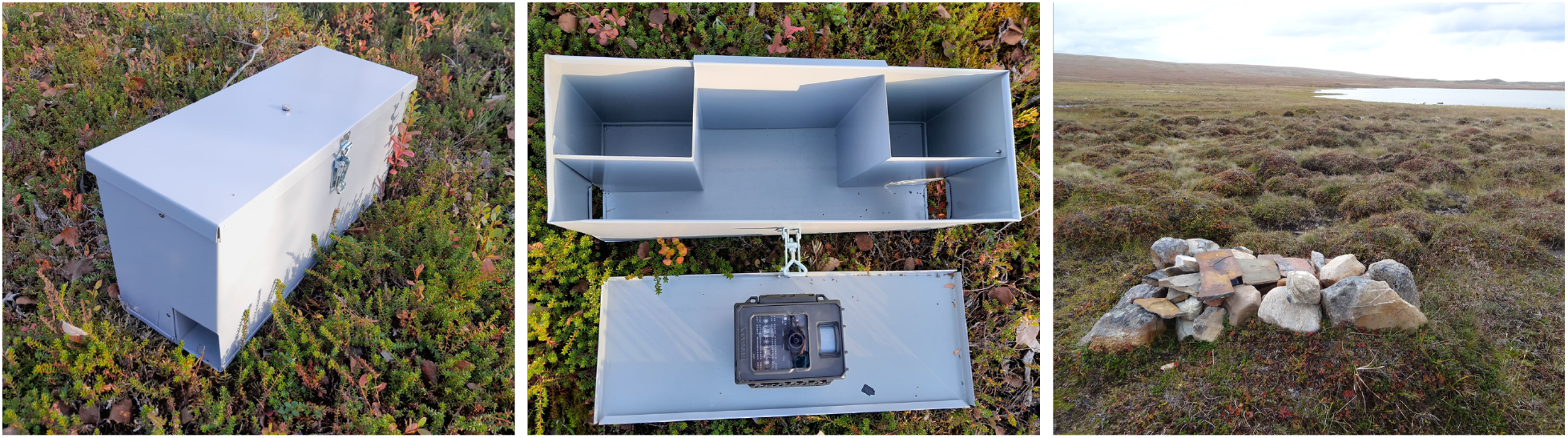
The camera trap developed by Soininen et al. (2015) consists of a Reconyx camera that is mounted on the ceiling of a metal box that can be entered by small mammals. The camera traps are protected with stones when placed in the field.

**Figure 4:**
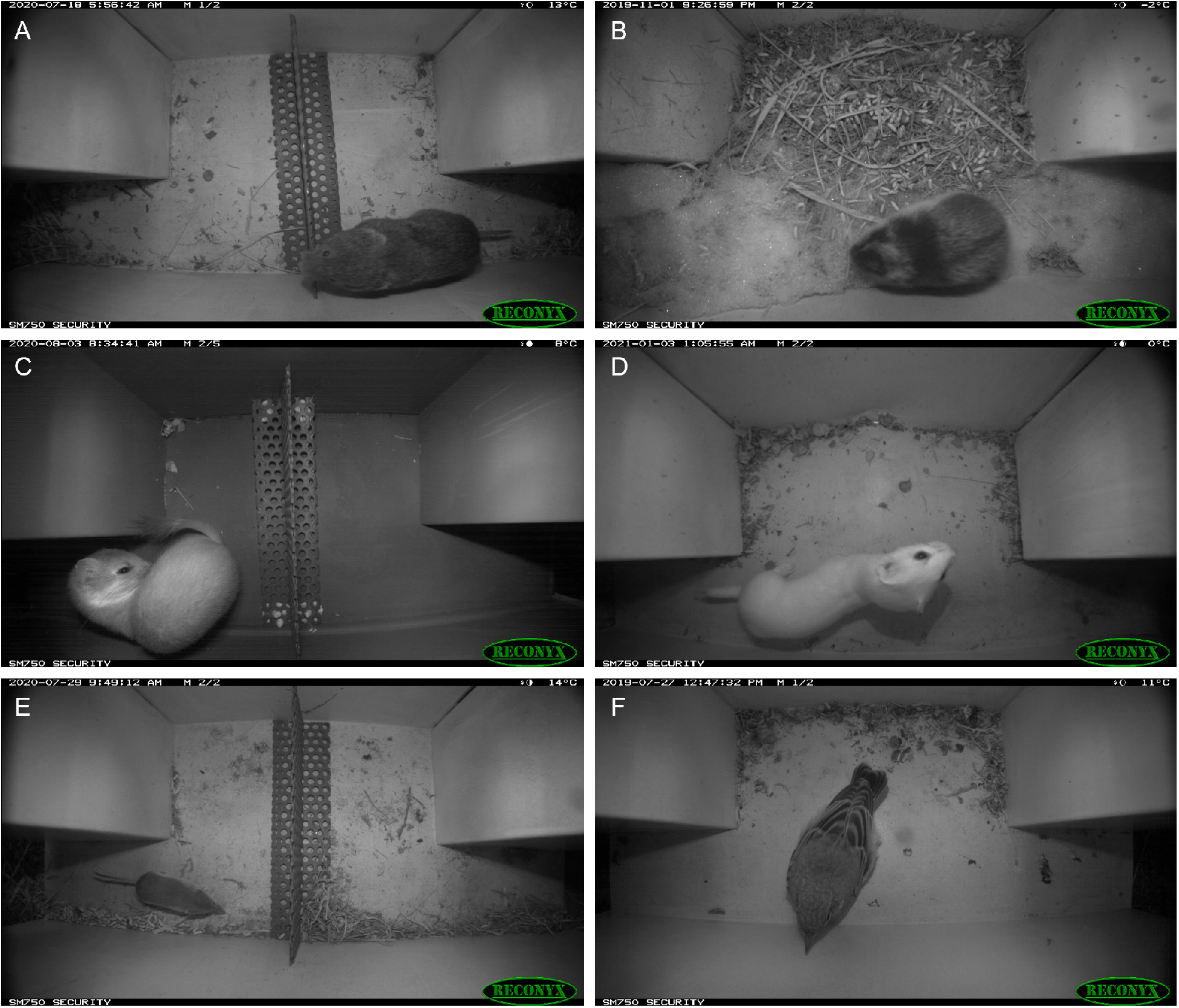
Example images of a vole (A), a lemming (B), a stoat (C), a least weasel (D), a shrew (E), and a bird (F).

In summer 2020, the long-term monitoring project was extended with 72 new camera trapping sites Varanger peninsula. These sites were established in the same two study areas as before (Komagdalen and Vestre Jakobselv) but in two new habitats (heath and meadow). We used a similar camera trap set up as for the initial traps, but deployed a new camera model (Customized from Reconyx Hyperfire II, Reconyx Inc., Holmen, WI, USA) inside the boxes, and the floor of the boxes was darker. Since the trigger speed of the new camera model was slower, which would lead to bad quality images if an animal runs through the camera box, a low metal barrier was installed in the middle of the box to slow down the animals. This made it possible to test if the classification model can be transferred to new camera trap sites. Furthermore, we could test if the camera model or modifications to the camera boxes have an effect on the performance of a model that was trained on slightly different images (Figure 4 A, C and E show boxes with the metal barrier).

### 3.2. Model training

To develop the classification model, we used the images taken between summer 2014 and summer 2020 in hummock tundra and snowbed habitat on the Varanger peninsula (original camera traps). Since our data set was unbalanced with a lot of empty and vole images but fewer images of stoats, least weasels and birds, we included images from other smaller small mammal camera trapping projects across Norway to increase the number of images of rare species. Furthermore, including more localities might potentially increase the transferability of the model to new camera trap sites. These camera traps are located in Porsanger, Kirkesdalen, Håkoya and Valdres with 3-15 camera trap sites per locality (Figure 2).

We selected 55922 images for model development from all available images taken between summer 2014 and summer 2020. The images were sorted in 6 animal classes (voles, lemmings, shrews, least weasels, stoats and birds), empty if there was no animal on the image and bad quality if it was not possible to decide whether the image was empty or not. Bad quality images are for example blurry images, images from boxes full of snow, water or vegetation or images of humans or landscapes taken when the camera was set up. All training images were selected manually and only images that could be easily sorted in one of the categories were included the training data set. We tried to create a balanced training data set by selecting images from all camera traps and by selecting a similar number of empty images and images of abundant species (voles, lemmings and shrews). We did not include difficult images since the quality of the training data set is important for the accuracy and the reliability of a neural network (Kavzoglu, 2009).

The selected 55922 images were then split in a training and a validation data set by selecting 350 (or less for rare species) random images per class for validation during the training process. In addition, we selected 4425 test images from the remaining images taken between summer 2014 and summer 2020 for testing the model after the training process was finished (Table 1).

**Table 1:** Number of training images, validation images used for model validation during model training and test images used for external model validation after training was finished as well as number of new images selected from the images taken between summer 2020 and summer 2021 for model retraining.

| Class | Number of<br>training images | Number of<br>validation images | Number of test<br>images | Number of new<br>training images<br>for model<br>retraining |
| --- | --- | --- | --- | --- |
| Bad<br>quality | 6453 | 350 | 748 | 306 |
| Bird | 3382 | 144 | 75 | 195 |
| Empty | 9444 | 350 | 1081 | 533 |
| Least<br>weasel | 1725 | 74 | 24 | 424 |
| Lemming | 9449 | 350 | 629 | 449 |
| Shrew | 9265 | 350 | 639 | 416 |
| Stoat | 4026 | 318 | 191 | 425 |
| Vole | 9894 | 350 | 1038 | 528 |
| TOTAL | 53636 | 2286 | 4425 | 3276 |

We trained a deep neural network in R (R Core Team, 2022) using KerasR (Chollet et al., 2017), an interface to TensorFlow (Abadi et al., 2015) for R. We used the ResNet50 architecture (He et al., 2016) and trained a model from scratch for 55 epochs with a one-cycle learning rate policy with a minimum learning rate of 0.000001 and a maximum learning rate of 0.001 (Smith, 2018). All images were resized to 224 x 224 pixels previous to training and image augmentation (shifts, horizontal flips, rotations, zooms and shears) were applied to expand the training data set.

We evaluated the model performance on the test data set by calculating model accuracy as the number of correct predictions divided by the number of all images as well as precision, recall and F1 score for each class:

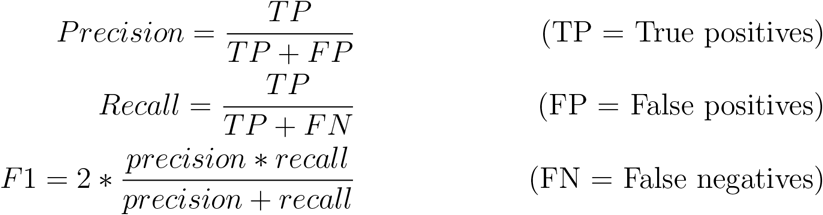

We also summarize true positives, false positives and false negatives in a confusion matrix created using the R package caret (Kuhn, 2008).

### 3.3. Data processing and semi-automatic workflow

To demonstrate the workflow for processing camera trap images and to test the transferability of the model to new sites and camera set ups, we used all images taken on the Varanger peninsula between summer 2020 and summer 2021 (including images from the original and the new camera traps).

After the images were downloaded from SD-cards in the field, we pre-processed the images by extracting meta data such as date, time and temperature from the images. Then, all images were renamed with a unique name including camera-site-id and the date when the image was taken. We classified all images automatically using our model and extracted the class with the highest confidence from the model output, which was used as the automatic image label.

We then quality checked the automatic image labels from the model in three levels: First, we calculated overall model accuracy for each camera trap type (i.e. original camera traps in hummock tundra and snowbed habitat and new camera traps in heath and meadow habitat). We did this by selecting 1000 random images per camera trap type, labeled them manually, and compared manual and automatic image labels. Second, to evaluate model performance for each class, we selected additional 200 random images per class and camera type and labeled them manually. We then calculated precision, recall and F1 score for each camera trap type using the 1000 random images and the 200 images per class. Including 200 images of each class increased the proportion of rare classes and thus, model performance would be overestimated for rare classes and underestimated for abundant classes. Therefore, the number of true positives, false positives and false negatives was corrected for the proportion of each class in the complete data set (see Appendix A.1). We also visualized these results in form of a confusion matrices (without correction) for each camera trap type using the R package caret (Kuhn, 2008). Third, to determine the confidence level above which the automatic image labels can be accepted, we selected 200 images per confidence class (0-0.1, 0.1-0.2, …, 0.9-1.0) and per camera trap type for calculating accuracy for each confidence class.

After the quality check, we decided to improve model performance by retraining the model with some of the new images (see section 3.4 for model retraining). We re-classified all images with the retrained model and repeated the quality check. Based on the different quality measures, we decided which automatic image labels we will accept and reviewed the remaining images manually.

### 3.4. Model updating

From all images taken in between summer 2020 and summer 2021 we added 3276 new images to the training data set. When selecting the new training images, we tried to select the same number of images from all camera trap sites and classes. Since images with stoats, least weasels and birds were underrepresented in the original training data set, we selected almost all images of these classes. If we selected many images from one site, we also selected some extra empty images, as well as images of other classes if available, from the same site and taken during the same period in order to prevent the model from associating the image background with a certain class. We did not take this into account when selecting the original training images and therefore also added some images from the previous years to the training data set. We then retrained the neural network as described before.

## 4. Results of the case study

### 4.1. Model training

Prediction accuracy of the small mammal classification model after training for 55 epochs evaluated on the test data set was 98.3%. 97.3% of the test images were classified with a confidence above 0.9 and 99.3% of these images were labeled correctly.

The model performed well for the classes with many training images (F1 score between 0.98 and 0.99), whereas model performance was poorer for the classes with less training images (stoat, least weasel and bird, Table 2, Figure 5).

**Table 2:** Precision, recall and F1 score for the 8 classes in the test data set predicted using the original and the retrained classification model. The test data set includes images taken between summer 2014 and summer 2020 with the original camera traps.

| Class | Original model |  |  | Retrained model |  |  |
| --- | --- | --- | --- | --- | --- | --- |
|  | Precision | Recall | F1 | Precision | Recall | F1 |
| Bad quality | 0.98 | 1 | 0.99 | 0.98 | 1 | 0.99 |
| Bird | 0.87 | 1 | 0.93 | 0.91 | 1 | 0.96 |
| Empty | 0.97 | 0.99 | 0.98 | 0.97 | 0.99 | 0.98 |
| Least weasel | 0.92 | 1 | 0.96 | 0.79 | 0.96 | 0.87 |
| Lemming | 1 | 0.99 | 0.99 | 1 | 0.98 | 0.99 |
| Shrew | 0.99 | 0.97 | 0.98 | 0.99 | 0.97 | 0.98 |
| Stoat | 0.99 | 0.97 | 0.98 | 0.99 | 0.95 | 0.97 |
| Vole | 0.99 | 0.97 | 0.98 | 0.99 | 0.98 | 0.99 |

**Figure 5:**
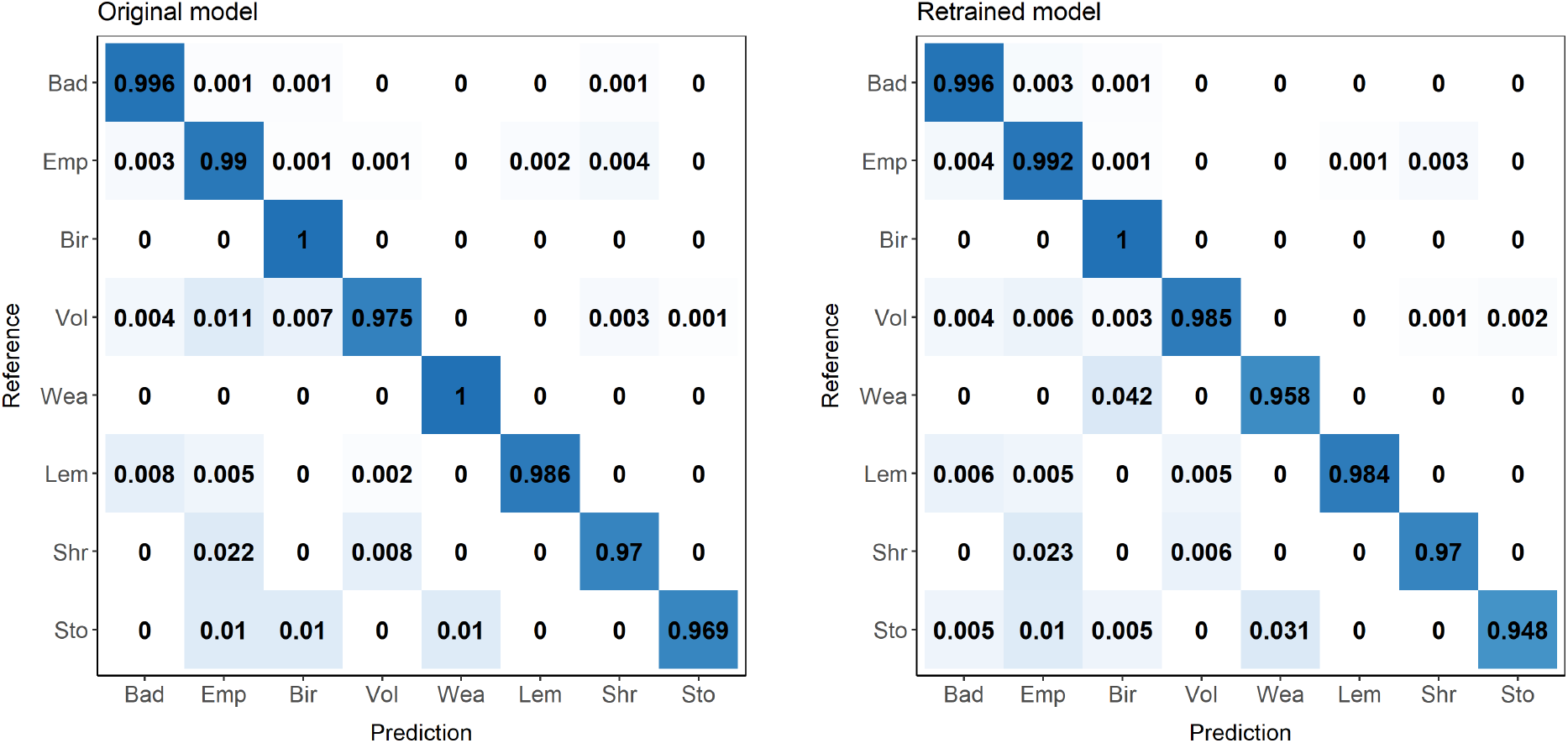
Confusion matrix (percentage of correct labels for each class) showing the performance of the original and the retrained classification model on the test data set. The test data set includes images taken between summer 2014 and summer 2020 with the original camera traps. (Bad = Bad quality, Emp = Empty, Bir = Bird, Vol = Vole, Wea = Least weasel, Lem = Lemming, Shr = Shrew, Sto = Stoat

### 4.2. Classification and quality check of new images with the original model

Between summer 2020 and summer 2021, the original camera traps in hummock tundra and snowbed habitat took 368735 images and the newly established camera traps in meadow and heath habitat took 70279 images. These images were automatically classified with our small mammal classification model.

The quality check showed that overall prediction accuracy was 95.3 % for the original camera traps and 90.7 % for the new camera traps (calculated using 1000 manually labeled images per camera trap type). Prediction accuracy increased with the model confidence (Figure 6). 88.2 % of the images from the original camera traps and 73.7 % of the images from the new camera traps were classified with a confidence above 0.9.

**Figure 6:**
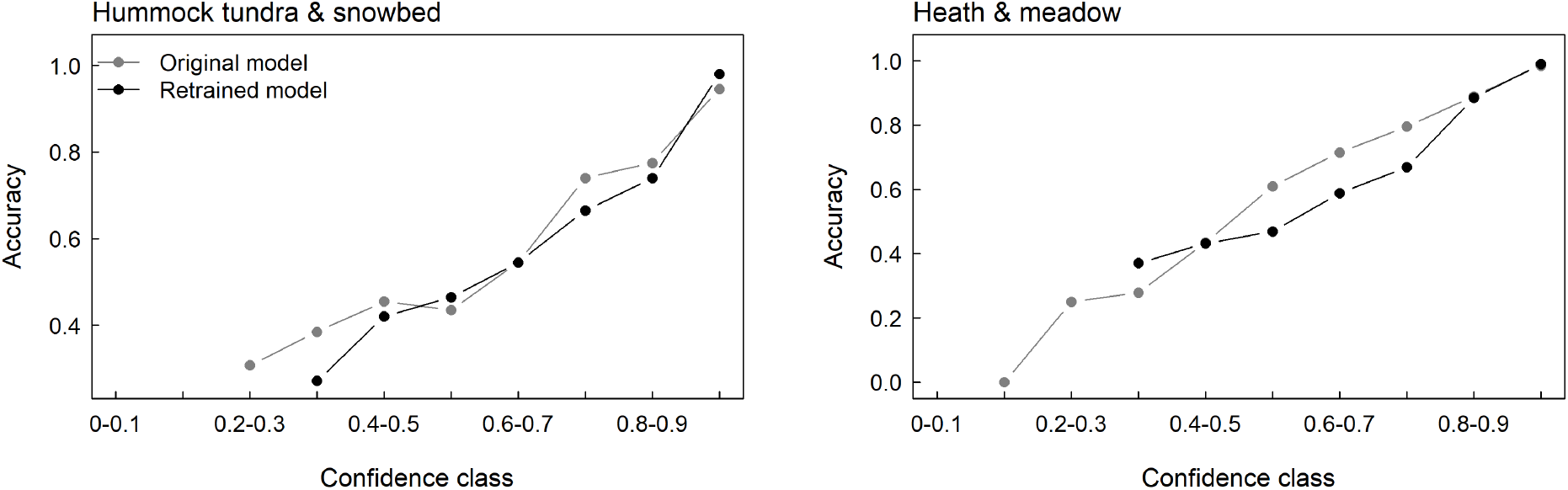
Prediction accuracy of images that were classified with a confidence between 0 and 0.1, between 0.1 and 0.2, …, and between 0.9 and 1.0 with the original and the retrained model. Prediction accuracy was calculated using 200 randomly selected and manually labeled images per camera trap type (original camera traps in hummock tundra and snowbed habitat and new camera traps in heath and meadow habitat) and confidence class.

### 4.3. Model retraining

After retraining the original small mammal classification model with new images from 2020-2021, prediction accuracy evaluated on the test data set was 98.5 %. 97.9 % of the images were classified with a confidence above 0.9 and 99.2 % of these images were labeled correctly.

### 4.4. Classification and quality check of new images with the retrained model

After classification of the new images taken between summer 2020 and summer 2021 with the retrained model, overall prediction accuracy was 98.4 % for the original camera traps and 97.5 % for the new camera traps. Prediction accuracy per confidence class was similar for both models and prediction accuracy of images classified with a confidence above 0.9 was 98 % for the new camera traps and 99 % for the original camera traps (Figure 6). However, retraining the model increased the percentage of images classified with a confidence above 0.9 for both camera trap types. 97.9 % of the images from the original camera traps and 94.9 % of the images from the new camera traps were classified with a confidence above 0.9 with the retrained model.

Retraining the model with images from 2020-2021 also improved model precision and recall of all classes. While model performance of the original model was poor for some classes, it was acceptable for the updated model with F1 scores above 0.8 for all classes (Table 3 and Figure 7).

**Table 3:** Precision, recall and F1 score for the 8 classes in the new data sets from 2020-2021 predicted using the original and the retrained classification model (bold numbers). The new data sets include the images taken between summer 2020 and summer 2021 with the original camera traps in hummock tundra and snowbed habitat and the new camera traps in heath and meadow habitat. The values presented here were corrected for the proportion of each class in the complete data set. The correction and the uncorrcted values are shown in Appendix A.1 and A.2.

| Class | Hummock tundra & Snowbed |  |  | Heath & Meadow |  |  |
| --- | --- | --- | --- | --- | --- | --- |
|  | Precision | Recall | F1 | Precision | Recall | F1 |
| Bad quality | (0.69)<br>**0.89** | (0.86)<br>**0.79** | (0.77)<br>**0.83** | (0.83)<br>**0.93** | (0.84)<br>**0.96** | (0.84)<br>**0.95** |
| Bird | (0.47)<br>**0.89** | (0.82)<br>**0.87** | (0.60)<br>**0.88** | (0.17)<br>**0.90** | (0.14)<br>**0.98** | (0.15)<br>**0.94** |
| Empty | (0.99)<br>**0.99** | (0.95)<br>**0.99** | (0.97)<br>**0.99** | (0.95)<br>**0.99** | (0.96)<br>**0.99** | (0.96)<br>**0.99** |
| Least weasel | (0.39)<br>**0.87** | (0.98)<br>**1.00** | (0.56)<br>**0.93** | (0.10)<br>**0.72** | (0.96)<br>**1.00** | (0.18)<br>**0.84** |
| Lemming | (0.79)<br>**0.80** | (1.00)<br>**1.00** | (0.88)<br>**0.89** | (0.67)<br>**0.85** | (0.96)<br>**0.99** | (0.79)<br>**0.92** |
| Shrew | (0.56)<br>**0.89** | (0.77)<br>**0.86** | (0.65)<br>**0.87** | (0.81)<br>**0.97** | (0.71)<br>**0.97** | (0.75)<br>**0.97** |
| Stoat | (0.35)<br>**0.82** | (0.80)<br>**1.00** | (0.49)<br>**0.90** |  |  |  |
| Vole | (0.93)<br>**0.99** | (0.95)<br>**0.97** | (0.94)<br>**0.98** | (0.96)<br>**0.99** | (0.85)<br>**0.96** | (0.90)<br>**0.97** |

**Figure 7:**
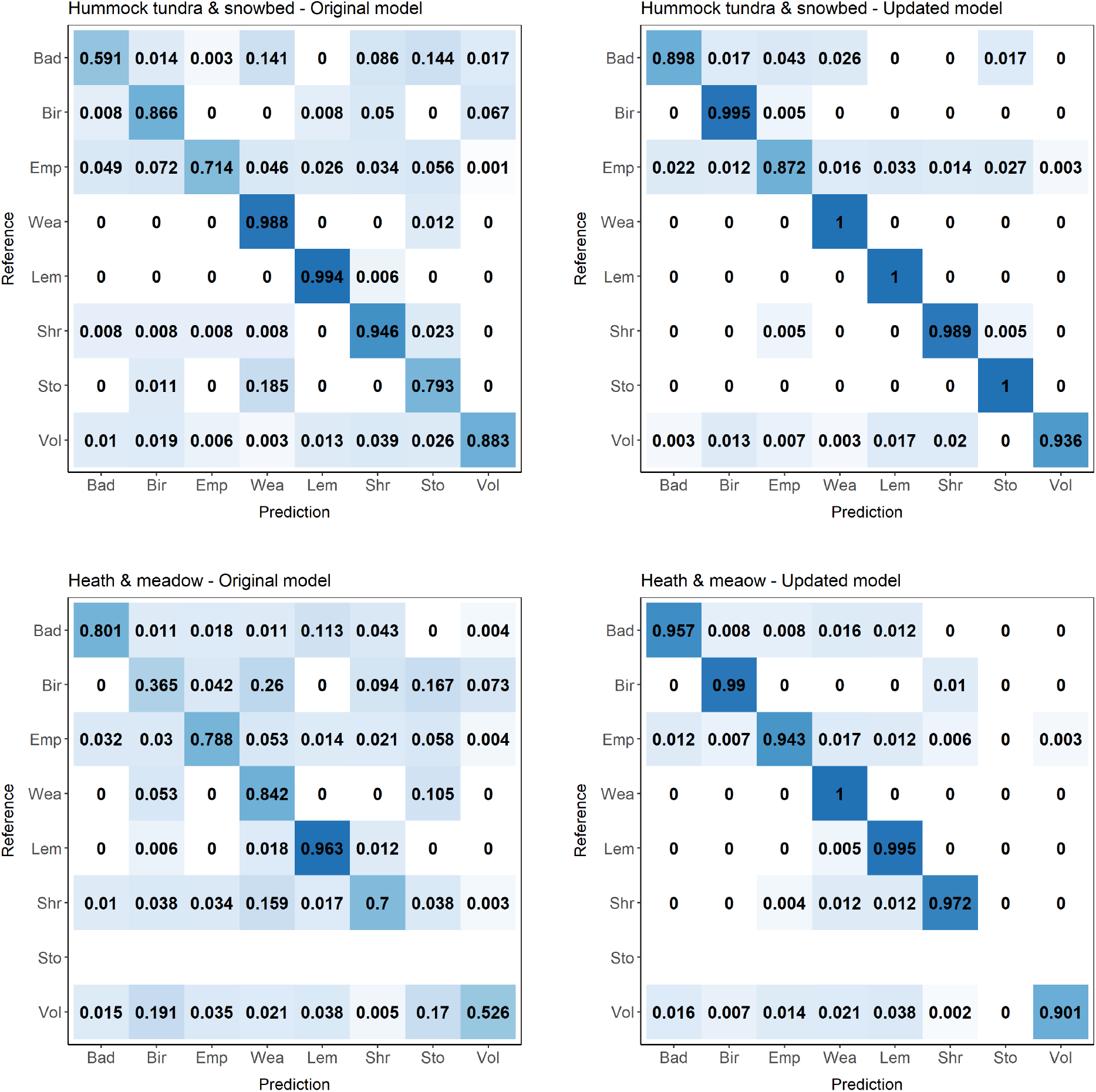
Confusion matrix (percentage of correct labels for each class) showing the performance of the original and the retrained classification model on new data sets from 2020-2021. The new data sets include the images taken between summer 2020 and summer 2021 with the original camera traps in hummock tundra and snowbed habitat and the new camera traps in heath and meadow habitat. (Bad = Bad quality, Emp = Empty, Bir = Bird, Vol = Vole, Wea = Least weasel, Lem = Lemming, Shr = Shrew, Sto = Stoat).

## 5. Discussion

We presented a semi-automatic workflow for processing camera trap images that we demonstrated using images from long-term monitoring of small mammals. We trained a classification model for the identification of small mammal species, since most available tools for automatic classification of camera trap images are designed for larger animals and were therefore not applicable for our data set. Our model had an accuracy of over 98 %, which is among the highest accuracies reached so far when using neural networks to classify camera trap images (Tabak et al., 2019). Our model also showed high precision and recall for all classes, although the training data set was somewhat unbalanced and relatively small (e.g. Norouzzadeh et al., 2018; Tabak et al., 2019). We trained the model, and constructed the entire workflow in R, to demonstrate that extensive machine learning knowledge or programming skills in python are not needed to implement automatic classification in camera trap studies. Instead, we believe that ecologists with an average knowledge of R are able to train their own model and set up their own workflow following our scripts. We illustrated the proposed workflow by processing new images, including images from new camera trap sites with modified camera traps. This is a typical scenario of a long-term monitoring project where new data is coming in periodically, new sites might be added and new camera models acquired. The flexibility, transparency, and required skill levels thus makes our workflow an especially useful tool for ecological long-term monitoring programs with large camera trap data sets.

Our workflow includes a relatively extensive quality check of automatic image labels. Such a check is crucial, as machine learning models usually decrease dramatically in accuracy when they are applied to new data (Schneider et al., 2020). Thus, automatically classifying images that are completely new to the model, for example images taken after the model has been trained, can result in poorer model performance and the verification of automatic image labels has been emphasized by several authors (Christin et al., 2021; Vélez et al., 2022). In our case study, the original model classified the new images with a prediction accuracy over 90 %, which is very high when a classification model is transferred to new data (Schneider et al., 2020). However, a drop of prediction accuracy from 98 % to 90 % from one year to the next can compromise the comparability of data from a long-term monitoring program. We could show that if retraining the model with new images is part of the workflow, model performance for new images can be increased strikingly (to 97.5 % for the the new camera traps and 98.4 % for the old camera traps). Even if overall prediction accuracy was high, the quality check revealed that images classified with a confidence above 0.9 had a prediction accuracy around 99 %, while images classified with a confidence between 0.8 and 0.9 had a prediction accuracy around 80 %. Therefore, we decided to accept all model labels with a confidence above 0.9 and labeled the remaining images manually. Furthermore, we also discovered differences in model performance for the difference classes, i.e. our model performed poorly for stoats and least weasels. Since these are target species of the monitoring, we also labeled all images of these classes manually. Another reason for an extensive quality check is that image data sets usually contain some images that are difficult for a classification model. For example, our data set contained one box with a dead lemming. Images from this box were all labeled as ‘lemming’ although the correct label would be ‘empty’. Furthermore, the boxes often get filled with snow during winter, and sometimes the snow patches have similar shapes as mustelids and are therefore labeled as ‘stoat’ or ‘least weasel’. We also found new a species (mink) on the some images from the new camera traps. If enough images of all classes and confidence-levels are reviewed in a quality check, these issues are likely to be discovered and can then be corrected.

Our workflow minimized the effort of human labeling through retraining the model with new images. Retraining the model with 3276 out of over 400.000 new images was enough to increase model accuracy as well as the percentage of images that were labeled with high confidence. This meant that only around 10.000 images were classified with a confidence below 0.9 and had to be reviewed manually, compared to around 60.000 images without retraining the model. We thus saved about 70 hours of manual image labeling when estimating that a human will label about 700 images per hour. Besides saving a lot of time, reducing the number of manually labeled images might also increase the accuracy of the data set. Human accuracy is likely to decrease if a large data set has to be reviewed because humans might be more prone to make mistakes when looking at images for hours and days. Furthermore, a small data set can often be labeled by experts whereas the help from less experienced assistants, who often obtain lower accuracies, is needed for labeling a large data set. For example, the accuracy of volunteers labeling a large data set was 96.6 % (Norouzzadeh et al., 2018) while we assume an accuracy of around 99 % for our data set. Therefore, we think an extensive quality check to identify images for which the model performs poorly and correcting the labels of these image as well as possibly retraining the model with new images are important steps of the workflow when a model is applied to new data. This will often be the case when camera traps are used long-term monitoring programs. Typically, a model is trained on images gathered over several years and will then be used to automatically classify images from the following years.

Our workflow was developed for images from small mammal camera boxes which are relatively easy for model development and transferability since all images have a similar background and usually contain only one individual. Other small mammal camera trap studies can directly implement our workflow by using the provided R scripts and instructions (https://github.com/hannaboe/camera_trap_workflow). If the same or similar species are target as in our data set (voles, lemmings, mustelids, shrews and small birds), our model can be used for automatic classification. We also provide our training data set, so the model can be retrained with images from other localities to increase model performance. If species composition differs from our data set, a new model could be trained using transfer learning with our model as a base-model. With transfer learning, good model performance can be reached even if the training data set is small (Weiss et al., 2016). Several studies successfully trained models with around 1000 images per class when using transfer learning (Shahinfar et al., 2020; Ferreira et al., 2020). Thus, new projects that are building on our model do not have to put a lot of effort in creating a large training data set to train their own models.

To our knowledge, our workflow is the first approach that gathers and streamlines all steps of a processing pipeline for camera trap images using automatic classification within one platform. Thus, it is not limited to small mammal camera trap studies but can also be an example for developing a processing workflow for other camera trap studies. And although the workflow was developed to be used in combination with a classification model (models that assign a certain category to an image), we think the workflow can be adjusted to be used with object detection models (i.e. models that draw bounding boxes around objects on the image and assign categories to each box). Due to its flexibility, the workflow is especially well suited for long-term monitoring programs where methods often have to be adapted over time. Sites, camera types or species composition might change over time and machine learning models and even software have to be updated. We provide R scripts for all steps of the workflow which can easily be adjusted to fit the specific needs of a new study or changes in the monitoring design. In addition, we also provide R-shiny-apps, which are as flexible as R scripts, for the more time consuming steps of the work (quality check and correction of model labels). After the apps have been adjusted to the study, they can be used without knowledge of R and even volunteers can help with labeling images manually. Alternatively, other software tools can be used to perform the different steps, even though switching between tools may be problematic. Furthermore, tools with a graphical user interface are often less flexible and more difficult to adjust for ecologists than R scripts. Thus, although many tools for processing camera trap images have been developed in the last years, we anticipate that the workflow presented here can help other research programs to develop their own routines for processing camera trap images and to incorporate automatic classification in their workflows.

## 6. Acknowledgements

We thank Siw Killengreen and Ingrid Jensvoll for initiating the development of small mammal camera trapping at Varanger Peninsula, Jan-Erik Knutsen, Torunn Moe and Leif-Einar Støvern for help with field logistics, and everyone who has participated in camera trapping field work since 2014: Siw Killengreen, Ingrid Jensvoll, Leif Einar Støvern, Dorothee Ehrich, Francisco Javier Ancin Murguzur, Jonas Mölle, Mathias Leines Dahle, Kjerstin Mæland, Jarad Mellard, Mike Murphy, Jan-Erik Knutsen, and Julia Mikhailova. The camera trapping infrastructure has been funded by the Research Council of Norway (grant no 245638). The same grant, together with UiT-The Arctic University of Norway has provided salary funds. This study is a contribution from the Climate-ecological Observatory for Arctic Tundra. The the images were stored and processed on resources provided by Sigma2 - the National Infrastructure for High Performance Computing and Data Storage in Norway

## 7. Data accessibility

All R scripts together with a some example images and detailed instructions how to set up the workflow are available on GitHub (https://github.com/hannaboe/camera_trap_workflow). The small mammal classification model as well as the images used for training, validating and testing the model are available at https://doi.org/10.5281/zenodo.7142734.

## 8. Author contributions

Conceptualisation (EMS, HB, EFK, RAI), Data Curation (HB, EFK), Formal Analysis (HB), Funding Aquisition (RAI, EMS), Investigation (HB, EFK, RAI, EMS), Methodology (HB), Project Administration (HB, EMS), Resources (RAI, EMS), Software (HB), Validation (HB, EFK, RAI, EMS), Visualization (HB), Writing - original draft (HB), Writing - review and editing (HB, EFK, RAI, EMS).

## Appendix A. Supplementary information

### Appendix A.1. Correction of true positives, false positives and false negatives

We calculated precision, recall and F1 score using 200 images per class in addition to 1000 randomly selected images. This increased the proportion of rare classes and thus, model performance would be overestimated for rare classes and underestimated for abundant classes. Therefore, the number of true positives, false positives and false negatives were corrected for the proportion of each class in the complete data set before precision, recall and F1 score were calculated.

~~~
#### Calculation of precision, recall and F1 score
### with correction of the number of true positives, false positives and false
### negatives for the proportion of each class in the complete data set
# complete is the complete data set with automatic image labels
# qual_check is a subset that was manually annotated for a quality check
# guess1 is the automatic image label
# class_id is the manual image label
----
classes <- unique(complete$guess1) # all classes in the complete data set
## calculate proportion of images per class in complete and quality check data set
prop_complete <- c()
prop_qual_check <- c()
for (i in 1:length(classes)) {
   prop_complete[i] <- length(which(complete$guess1 == classes[i]))/nrow(complete)
   prop_qual_check[i] <- length(which(dat$guess1 == classes[i]))/nrow(dat)
} proportion <- data.frame(class_id = classes, prop_complete = prop_complete, prop_qual_check = prop_qual_check)
## correct true positives, false positives and
## calculate precision, recall and F1 score for each class
for (i in 1:length(classes)){
   ## proportion of class i in the two datasets
  prop_complete <- proportion$prop_complete[proportion$class_id == classes[i]]
  prop_qual_check <- proportion$prop_qual_check[proportion$class_id == classes[i]]
## calculate true posivies, false positives and false negatives
tp <- dat[dat$class_id == classes[i] & dat$guess1==classes[i],]
fp <- dat[dat$class_id != classes[i] & dat$guess1==classes[i],]
fn <- dat[dat$class_id == classes[i] & dat$guess1 != classes[i],]
## correct true positives and false positives
tp_corr <- nrow(tp)/prop_qual_check*prop_complete
fp_corr <- nrow(fp)/prop_qual_check*prop_complete
## correct fn
fn_corr <- c()
for (j in 1:length(classes)) {
   if (classes[j] %in% fn$guess1) {
     prop_complete2 <- proportion$prop_complete[proportion$class_id == classes[j]]
     prop_qual_check2 <- proportion$prop_qual_check[proportion$class_id == classes[j]]
     fn_corr[j] <- nrow(fn[fn$guess1 == classes[j],])/prop_qual_check2*prop_complete2
  }
 }
 fn_corr <- sum(fn_corr, na.rm = TRUE)
 ## calculate precision, recall and F1 score
 precision <- tp_corr/(tp_corr+fp_corr)
 recall <- tp_corr/(tp_corr+fn_corr)
 f1 <- 2*((precision*recall)/(precision+recall))
}
~~~

### Appendix A.2. Precision, Recall and F1 score without correction

**Table A.4:** Precision, recall and F1 score for the 8 classes in the new data sets from 2020-2021 predicted using the original and the retrained classification model (bold numbers). The new data sets include the images taken between summer 2020 and summer 2021 by the original camera traps in hummock tundra and snowbed habitat and the new camera traps in heath and meadow habitat. Precision, recall and F1 score were calculted using the R-package caret (Kuhn, 2008)

| Class | Hummock tundra & Snowbed |  |  | Heath & Meadow |  |  |
| --- | --- | --- | --- | --- | --- | --- |
|  | Precision | Recall | F1 | Precision | Recall | F1 |
| Bad quality | (0.70)<br>**0.88** | (0.59)<br>**0.90** | (0.64)<br>**0.89** | (0.83)<br>**0.93** | (0.80)<br>**0.96** | (0.81)<br>**0.94** |
| Bird | (0.48)<br>**0.89** | (0.87)<br>**0.99** | (0.61)<br>**0.94** | (0.17)<br>**0.90** | (0.36)<br>**0.99** | (0.23)<br>**0.94** |
| Empty | (1.00)<br>**0.99** | (0.71)<br>**0.87** | (0.83)<br>**0.93** | (0.95)<br>**0.99** | (0.79)<br>**0.94** | (0.86)<br>**0.97** |
| Least weasel | (0.40)<br>**0.87** | (0.99)<br>**1.00** | (0.56)<br>**0.93** | (0.10)<br>**0.72** | (0.84)<br>**1.00** | (0.18)<br>**0.84** |
| Lemming | (0.81)<br>**0.80** | (0.99)<br>**1.00** | (0.89)<br>**0.89** | (0.67)<br>**0.85** | (0.96)<br>**0.99** | (0.79)<br>**0.92** |
| Shrew | (0.57)<br>**0.89** | (0.95)<br>**0.99** | (0.72)<br>**0.94** | (0.81)<br>**0.97** | (0.7)<br>**0.97** | (0.75)<br>**0.97** |
| Stoat | (0.35)<br>**0.82** | (0.80)<br>**1.00** | (0.49)<br>**0.90** |  |  |  |
| Vole | (0.95)<br>**0.99** | (0.88)<br>**0.94** | (0.91)<br>**0.96** | (0.96)<br>**0.99** | (0.52)<br>**0.90** | (0.68)<br>**0.94** |

## References

Abadi, M., Agarwal, A., Barham, P., Brevdo, E., Chen, Z., Citro, C., Corrado, G.S., Davis, A., Dean, J., Devin, M., Ghemawat, S., Goodfellow, I., Harp, A., Irving, G., Isard, M., Jia, Y., Jozefowicz, R., Kaiser, L., Kudlur, M., Levenberg, J., Mané, D., Monga, R., Moore, S., Murray, D., Olah, C., Schuster, M., Shlens, J., Steiner, B., Sutskever, I., Talwar, K., Tucker, P., Vanhoucke, V., Vasudevan, V., Viégas, F., Vinyals, O., Warden, P., Wattenberg, M., Wicke, M., Yu, Y., Zheng, X., 2015. TensorFlow: Large-scale machine learning on heterogeneous systems. URL: https://www.tensorflow.org/. software available from tensorflow.org.

Beery, S., Morris, D., Yang, S., Simon, M., Norouzzadeh, A., Joshi, N., 2019. Efficient pipeline for automating species id in new camera trap projects. Biodiversity Information Science and Standards 3, e37222. URL: doi:10.3897/biss.3.37222, arXiv:https://doi.org/10.3897/biss.3.37222.

Burton, A.C., Neilson, E., Moreira, D., Ladle, A., Steenweg, R., Fisher, J.T., Bayne, E., Boutin, S., 2015. REVIEW: Wildlife camera trapping: a review and recommendations for linking surveys to ecological processes. Journal of Applied Ecology 52, 675–685. https://onlinelibrary.wiley.com/doi/abs/10.1111/1365-2664.12432 https://besjournals.onlinelibrary.wiley.com/doi/10.1111/1365-2664.12432, doi:10.1111/1365-2664.12432.

Chollet, F., Allaire, J., et al., 2017. R interface to keras. https://github.com/rstudio/keras.

Christin, S., Hervet, E., Lecomte, N., 2019. Applications for deep learning in ecology. Methods in Ecology and Evolution 10, 1632–1644. URL: https://onlinelibrary.wiley.com/doi/abs/10.1111/2041-210X.13256, doi:10.1111/2041-210X.13256.

Christin, S., Hervet, E., Lecomte, N., 2021. Going further with model verification and deep learning. Methods in Ecology and Evolution 12, 130–134. URL: https://onlinelibrary.wiley.com/doi/full/10.1111/2041-210X.13494 https://onlinelibrary.wiley.com/doi/abs/10.1111/2041-210X.13494 https://besjournals.onlinelibrary.wiley.com/doi/10.1111/2041-210X.13494, doi:10.1111/2041-210X.13494.

Farley, S.S., Dawson, A., Goring, S.J., Williams, J.W., 2018. Situating Ecology as a Big-Data Science: Current Advances, Challenges, and Solutions. BioScience 68, 563–576. URL: https://academic.oup.com/bioscience/article/68/8/563/5049569, doi:10.1093/BIOSCI/BIY068.

Ferreira, A.C., Silva, L.R., Renna, F., Brandl, H.B., Renoult, J.P., Farine, D.R., Covas, R., Doutrelant, C., 2020. Deep learning-based methods for individual recognition in small birds. Methods in Ecology and Evolution 11, 1072–1085. URL: https://onlinelibrary.wiley.com/doi/full/10.1111/2041-210X.13436 https://onlinelibrary.wiley.com/doi/abs/10.1111/2041-210X.13436 https://besjournals.onlinelibrary.wiley.com/doi/10.1111/2041-210X.13436, doi:10.1111/2041-210X.13436.

Glover-Kapfer, P., Soto-Navarro, C.A., Wearn, O.R., Carolina Soto-Navarro, C.A., Environment, U., 2019. Camera-trapping version 3.0: current constraints and future priorities for development. Remote Sensing in Ecology and Conservation 5, 209–223. URL: https://onlinelibrary.wiley.com/doi/full/10.1002/rse2.106 https://onlinelibrary.wiley.com/doi/abs/10.1002/rse2.106 https://zslpublications.onlinelibrary.wiley.com/doi/10.1002/rse2.106, doi:10.1002/RSE2.106.

Greenberg, S., Automated Image Recognition for Wildlife Camera Traps: Making it Work for You 1 URL: https://speciesclassification.westus2.cloudapp.azure.com/.

He, K., Zhang, X., Ren, S., Sun, J., 2016. Identity mappings in deep residual networks, in: Leibe, B., Matas, J., Sebe, N., Welling, M. (Eds.), Computer Vision – ECCV 2016, Springer International Publishing, Cham. pp. 630–645.

Ims, R., Jepsen, Jane, U., Stien, A., Yoccoz, Nigel, G., 2013. Science plan for coat: Climate-ecological observatory for arctic tundra. Fram Centre Report Series 1.

Kavzoglu, T., 2009. Increasing the accuracy of neural network classification using refined training data. Environmental Modelling and Software 24, 850–858. doi:10.1016/J.ENVSUFT.2008.11.012.

Kellenberger, B., Tuia, D., Morris, D., 2020. AIDE: Accelerating image-based ecological surveys with interactive machine learning. Methods in Ecology and Evolution 11, 1716–1727. doi:10.1111/2041-210X.13489.

Krizhevsky, A., Sutskever, I., Hinton, G.E., 2012. Imagenet classification with deep convolutional neural networks, in: Pereira, F., Burges, C.J.C., Bottou, L., Weinberger, K.Q. (Eds.), Advances in Neural Information Processing Systems, Curran Associates, Inc. URL: https://proceedings.neurips.cc/paper/2012/file/c399862d3b9d6b76c8436e924a68c45b-Paper.pdf.

Kuhn, M., 2008. Building predictive models in r using the caret package. Journal of Statistical Software, Articles 28, 1–26. URL: https://www.jstatsoft.org/v028/i05, doi:10.18637/jss.v028.i05.

Lai, J., Lortie, C.J., Muenchen, R.A., Yang, J., Ma, K., 2019. Evaluating the popularity of R in ecology. Ecosphere 10, e02567. URL: https://onlinelibrary.wiley.com/doi/full/10.1002/ecs2.2567 https://onlinelibrary.wiley.com/doi/abs/10.1002/ecs2.2567 https://esajournals.onlinelibrary.wiley.com/doi/10.1002/ecs2.2567, doi:10.1002/ECS2.2567.

Mölle, J.P., Kleiven, E.F., Ims, R.A., Soininen, E.M., 2022. Using subnivean camera traps to study Arctic small mammal community dynamics during winter. Arctic Science 8, 183–199. URL: https://cdnsciencepub.com/doi/full/10.1139/as-2021-0006, doi:10.1139/AS-2021-0006/ASSET/IMAGES/AS-2021-0006TAB2.GIF.

Norouzzadeh, M.S., Nguyen, A., Kosmala, M., Swanson, A., Palmer, M.S., Packer, C., Clune, J., 2018. Automatically identifying, counting, and describing wild animals in camera-trap images with deep learning. Proceedings of the National Academy of Sciences 115, E5716–E5725. URL: https://www.pnas.org/content/115/25/E5716, doi: 10.1073/PNAS.1719367115.

R Core Team, 2022. R: A Language and Environment for Statistical Computing. R Foundation for Statistical Computing. Vienna, Austria. URL: https://www.R-project.org/.

Schneider, S., Greenberg, S., Taylor, G.W., Kremer, S.C., 2020. Three critical factors affecting automated image species recognition performance for camera traps. Ecology and Evolution 10, 3503–3517. URL:https://onlinelibrary.wiley.com/doi/full/10.1002/ece3.6147 https://onlinelibrary.wiley.com/doi/abs/10.1002/ece3.6147 https://onlinelibrary.wiley.com/doi/10.1002/ece3.6147, doi:10.1002/ECE3.6147.

Shahinfar, S., Meek, P., Falzon, G., 2020. “How many images do I need?” Understanding how sample size per class affects deep learning model performance metrics for balanced designs in autonomous wildlife monitoring. Ecological Informatics 57, 101085. doi:10.1016/j.ecoinf.2020.101085.

Smith, L.N., 2018. A DISCIPLINED APPROACH TO NEURAL NETWORK HYPER-PARAMETERS: PART 1 – LEARNING RATE, BATCH SIZE, MOMENTUM, AND WEIGHT DECAY. URL: https://github.com/lnsmith54/hyperParam1., arXiv:1803.09820.

Soininen, E.M., Jensvoll, I., Killengreen, S.T., Ims, R.A., 2015. Under the snow: a new camera trap opens the white box of subnivean ecology. Remote Sensing in Ecology and Conservation 1, 29–38. URL: https://onlinelibrary.wiley.com/doi/full/10.1002/rse2.2 https://onlinelibrary.wiley.com/doi/abs/10.1002/rse2.2 https://zslpublications.onlinelibrary.wiley.com/doi/10.1002/rse2.2, doi:10.1002/RSE2.2.

Sokolova, M., Lapalme, G., 2009. A systematic analysis of performance measures for classification tasks. Information Processing Management 45, 427–437. doi:10.1016/J.IPM.2009.03.002.

Steenweg, R., Hebblewhite, M., Kays, R., Ahumada, J., Fisher, J.T., Burton, C., Townsend, S.E., Carbone, C., Rowcliffe, J.M., Whittington, J., Brodie, J., Royle, J.A., Switalski, A., Clevenger, A.P., Heim, N., Rich, L.N., 2017. Scaling-up camera traps: monitoring the planet’s biodiversity with networks of remote sensors. Frontiers in Ecology and the Environment 15, 26–34. URL: https://onlinelibrary.wiley.com/doi/full/10.1002/fee.1448 https://onlinelibrary.wiley.com/doi/abs/10.1002/fee.1448 https://esajournals.onlinelibrary.wiley.com/doi/10.1002/fee.1448, doi:10.1002/FEE.1448.

Tabak, M.A., Norouzzadeh, M.S., Wolfson, D.W., Newton, E.J., Boughton, R.K., Ivan, J.S., Odell, E.A., Newkirk, E.S., Conrey, R.Y., Stenglein, J., Iannarilli, F., Erb, J., Brook, R.K., Davis, A.J., Lewis, J., Walsh, D.P., Beasley, J.C., VerCauteren, K.C., Clune, J., Miller, R.S., 2020. Improving the accessibility and transferability of machine learning algorithms for identification of animals in camera trap images: MLWIC2. Ecology and Evolution 10, 10374–10383. URL: https://onlinelibrary.wiley.com/doi/10.1002/ece3.6692, doi:10.1002/ece3.6692.

Tabak, M.A., Norouzzadeh, M.S., Wolfson, D.W., Sweeney, S.J., Vercauteren, K.C., Snow, N.P., Halseth, J.M., Di Salvo, P.A., Lewis, J.S., White, M.D., Teton, B., Beasley, J.C., Schlichting, P.E., Boughton, R.K., Wight, B., Newkirk, E.S., Ivan, J.S., Odell, E.A., Brook, R.K., Lukacs, P.M., Moeller, A.K., Mandeville, E.G., Clune, J., Miller, R.S., 2019. Machine learning to classify animal species in camera trap images: Applications in ecology. Methods in Ecology and Evolution 10, 585–590. URL: https://onlinelibrary.wiley.com/doi/abs/10.1111/2041-210X.13120, doi:10.1111/2041-210X.13120.

Vélez, J., Castiblanco-Camacho, P.J., Tabak, M.A., Chalmers, C., Fergus, P., Fieberg, J., 2022. Choosing an Appropriate Platform and Workflow for Processing Camera Trap Data using Artificial Intelligence URL: http://arxiv.org/abs/2202.02283, arXiv:2202.02283v1.

Weiss, K., Khoshgoftaar, T.M., Wang, D.D., 2016. A survey of transfer learning. Journal of Big Data 3, 1–40. URL: https://journalofbigdata.springeropen.com/articles/10.1186/s40537-016-0043-6, doi:10.1186/S40537-016-0043-6/TABLES/6.

Willi, M., Pitman, R.T., Cardoso, A.W., Locke, C., Swanson, A., Boyer, A., Veldthuis, M., Fortson, L., 2019. Identifying animal species in camera trap images using deep learning and citizen science. Methods in Ecology and Evolution 10, 80–91. doi:10.1111/2041-210X.13099.

Young, S., Rode-Margono, J., Amin, R., 2018. Software to facilitate and streamline camera trap data management: A review. Ecology and Evolution 8, 9947–9957. doi:10.1002/ECE3.4464.

Zualkernan, I., Dhou, S., Judas, J., Sajun, A.R., Gomez, B.R., Hussain, L.A., 2022. An IoT System Using Deep Learning to Classify Camera Trap Images on the Edge. Computers 2022, Vol. 11, Page 13 11, 13. URL: https://www.mdpi.com/2073-431X/11/1/13/htm https://www.mdpi.com/2073-431X/11/1/13, doi:10.3390/CUMPUTERS11010013.

